# Isolating the Direct Effects of Growth Hormone on Lifespan and Metabolism

**DOI:** 10.1101/2024.09.18.613718

**Authors:** Alexander Tate Lasher, Kaimao Liu, Michael Fitch, Liou Y. Sun

## Abstract

Prior studies show that disrupting somatotropic axis components extends laboratory mouse lifespan, but confounding effects of additional genes and hormones obscure the specific impact of growth hormone (GH) on longevity. We address this issue by using mice with a specific knockout of the GH gene, revealing that disrupting GH alone substantially increases lifespan. The longevity effects are accompanied by altered metabolic fuel utilization, directly linking GH action to aging mechanisms.

## Introduction

The somatotrophic axis, comprised of GH-releasing hormone (GHRH) and GH secreted from the hypothalamus or pituitary, respectively, is a powerful determinant of laboratory mouse longevity evidenced by the dramatic lifespan extensions that result from genetic interruption at any level of this axis in mice (Bartke, 2021; Bartke et al., 2013). This body of work suggests that the action of GH is a critical regulator of mammalian lifespan. A crucial limitation of these studies, however, is that mice typically treated as “GH-deficient” display defects in several other genes and hormones which leaves the direct contribution of GH unexplored. Ames dwarf and Snell dwarf mice, deficient for GH as well as prolactin and thyroid-stimulating hormone, were among the first mice with defective somatotrophic signaling found to be long-lived (Brown-Borg et al., 1996; Flurkey et al., 2001). Mutant mice lacking a functional GHRH-receptor or functional GHRH also cannot be considered true models of “isolated GH deficiency” as the extrapituitary effects of GHRH, which have gained appreciation as important physiological regulators (Granata, 2016), could contribute to the lifespan extension reported in these mice (Flurkey et al., 2001; Sun et al., 2013). Additionally, mice with a targeted disruption of the GH-receptor (GHR) gene display dramatically elevated levels of GH (Zhou et al., 1997).

These caveats in mouse models of somatotrophic interruption have proven to be biologically relevant. Calorie restriction, an intervention known to extend lifespan (Swindell, 2012), further enhances the longevity of Ames dwarf (Bartke et al., 2001) and GHRH deficient (Sun et al., 2013) mice but has no effect on the average life of GHR knockout mice (Bonkowski et al., 2006). Further, it has recently been shown that double mutant GHRH-KO x GHR-KO mice display significantly lower IGF-1, leptin, and adiponectin levels compared to WT, GHR-KO, or GHRH-KO counterparts (Icyuz et al., 2021). The divergent phenotypes of these models of somatotrophic disruption and differential response to lifespan-extending interventions clearly indicate they cannot be treated as analogous and bring the specific contribution of GH into question. To address this critical gap in knowledge, we carried out the first assessment (to our very best knowledge) of lifespan in mice with a targeted GH gene knockout in conjunction with metabolic assessment during adulthood.

## Results

GH knockout (KO) mice on a mixed C57BL/6N x C57BL/6J x BALB/cByJ genetic background maintained under specific pathogen-free conditions with *ad-lib* access to standard rodent diet (NIH-31) and water displayed a 21% extension in median lifespan over WT littermates (2.15 years vs. 2.60 years; Fig. 1a), a significant extension in life (p=0.0011, logrank test; table 1). Cox proportional hazard analysis revealed significantly lower hazard ratios (p=0.0012; table 1), and survival at the 75^th^ (p=0.0023) and 90^th^ (p=0.0303) percentiles were significantly higher in KO mice when assessed by quantile regression (Wang et al., 2004) (table 1). When sexes were analyzed separately, KO males displayed a 27% extension in median lifespan over WT males (1.96 years vs. 2.48 years; Fig. 1b-c), a significant lifespan extension (p=0.0380, logrank test; p=0.0427, Cox proportional hazard; table 1). KO females displayed a more modest, yet still statistically significant, 14% extension in lifespan (2.37 years vs 2.69 years; Fig. 1b, 1d) over WT females (p=0.0038, logrank test; p=0.0041 Cox proportional hazard; table 1). We did not detect any differences in the lifespans of males and females within the same genotype (table 1).

**Table 1.** Statistical analysis of lifespans.

| Sex | Group | Median Lifespan | Logrank p-value | Cox Proportional Hazard |  | Maximal Lifespan |  |
| --- | --- | --- | --- | --- | --- | --- | --- |
|  |  |  |  | Hazard Ratio | p-value | 75th p-value | 90th p-value |
| Pooled | WT (n=40) | 2.15 years | 0.0011 | 1 | 0.0012 | 0.0023 | 0.0303 |
|  | KO (n=43) | 2.60 years |  | 0.4699 |  |  |  |
| Male | WT (n=20) | 1.96 years | 0.0380 | 1 | 0.0427 | NA | NA |
|  | KO (n=22) | 2.48 years |  | 0.5230 |  |  |  |
| Female | WT (n=20) | 2.37 years | 0.0038 | 1 | 0.0041 | NA | NA |
|  | KO (n=21) | 2.69 years |  | 0.352 |  |  |  |
| Male WT (n=20) |  | 1.96 years | 0.36 | 1 | 0.417 | NA | NA |
| Female WT (n=20) |  | 2.37 years |  | 0.7671 |  |  |  |
| Male KO (n=22) |  | 2.48 years | 0.62 | 1 | 0.623 | NA | NA |
| Female KO (n=21) |  | 2.69 years |  | 0.8562 |  |  |  |

Note: Maximal lifespan was analyzed for pooled sexes as the low sample size that results from stratifying sexes prevented meaningful statistical comparisons.

**Figure 1.**
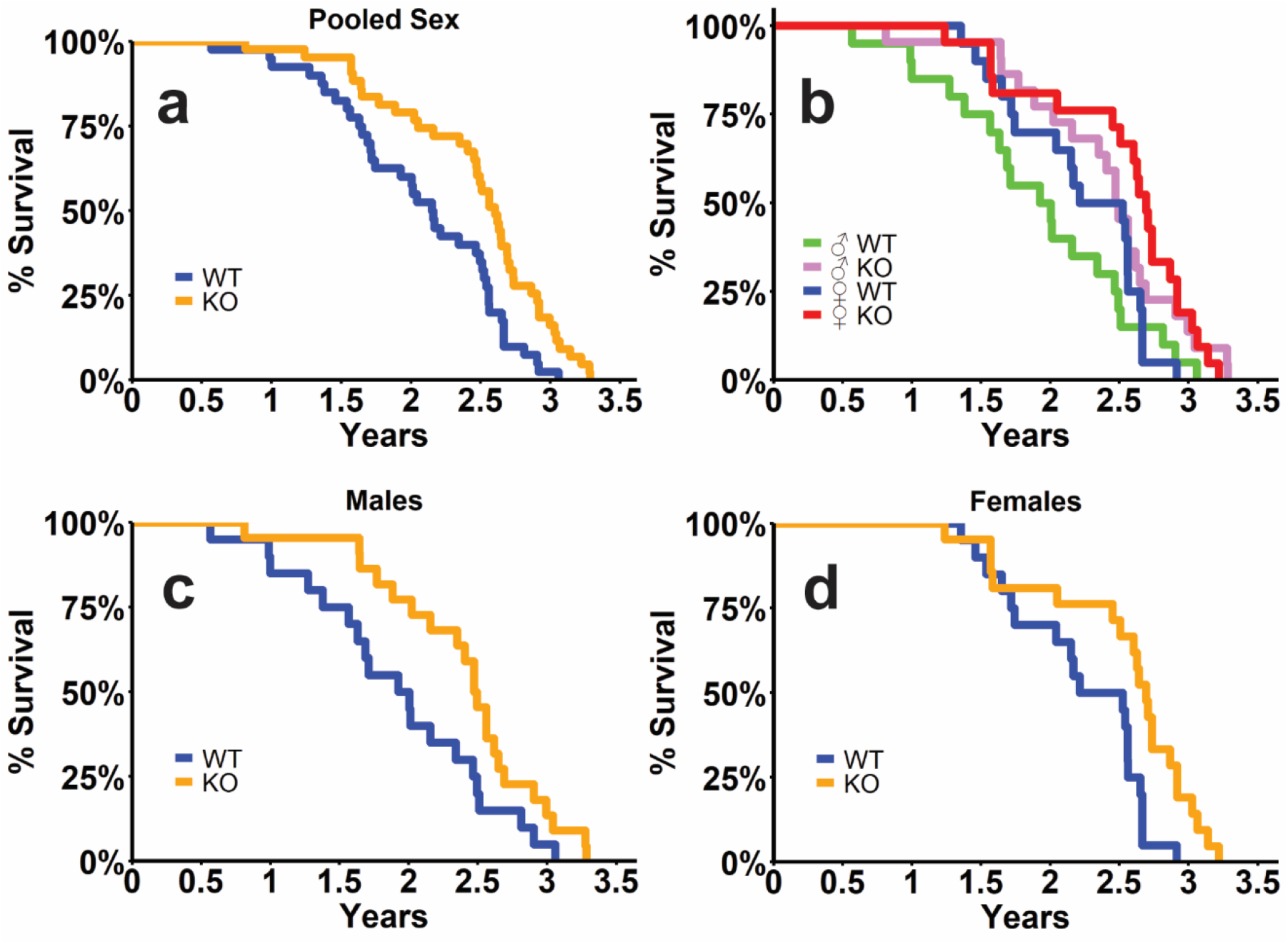
Isolated GH deficiency extends lifespan. Kaplan-Meier survival curves for pooled sexes (a), all groups (b), male mice (c) and female mice (d). Statistical analyses of lifespans are provided in table 1. WT = wild type; KO = GH knockout mice. N = 20 male WT; n = 22 male KO; n = 20 female WT; n = 21 female KO mice.

Physiological analysis of a separate cohort of mice showed reduced body weights and disproportionately higher body fat (Fig. 2a-d), consistent with previous reports (List et al., 2019), as well as lower food consumption and cage activity in KO mice (Supplemental Fig. S1a-d). Nutrient utilization and metabolic rate assessed by indirect calorimetry revealed significantly lower respiratory quotient (RQ) in male KOs during the light cycle during both light/dark cycles in female KOs (Fig. 2e, f) as well as lower O_2_ consumption and metabolic rates after accounting for bodyweight differences (Supplemental Fig. S1e-l). RQ varies inversely with fat oxidation (Zurlo et al., 1990), prompting us to explore fuel utilization more fully in these mice. Calculation of glucose or fat oxidation rates (GOx or FOx) as previously (Lasher & Sun, 2023) revealed marked reduction light cycle male KO GOx (Fig. 2g) and significant elevations in light cycle FOx (Fig. 2i). The reduced GOx (Fig. 2h) and elevated FOx (Fig. 2j) was observed during light/dark cycles for female KOs. These measures indicate that while fat utilization is certainly elevated in GH-deficient mice, consistent with typical interpretations of lower RQ (Icyuz et al., 2020; Icyuz et al., 2021), significant reductions in glucose utilization also exist in these KO mice indicating they rely on fat oxidation and less on glucose oxidation to meet energy demands. Further supporting the disrupted glucose metabolism in GH-deficient mice is the impaired glucose tolerance, evidenced by elevated glycemia in males and females following a glucose challenge (Fig. 2k, l). While the impaired glucose tolerance observed here is consistent with a previous report in this model (List et al., 2019), insulin sensitivity was enhanced only in male KOs and unchanged in female mice (Supplemental Fig. 1m, n).

**Figure 2.**
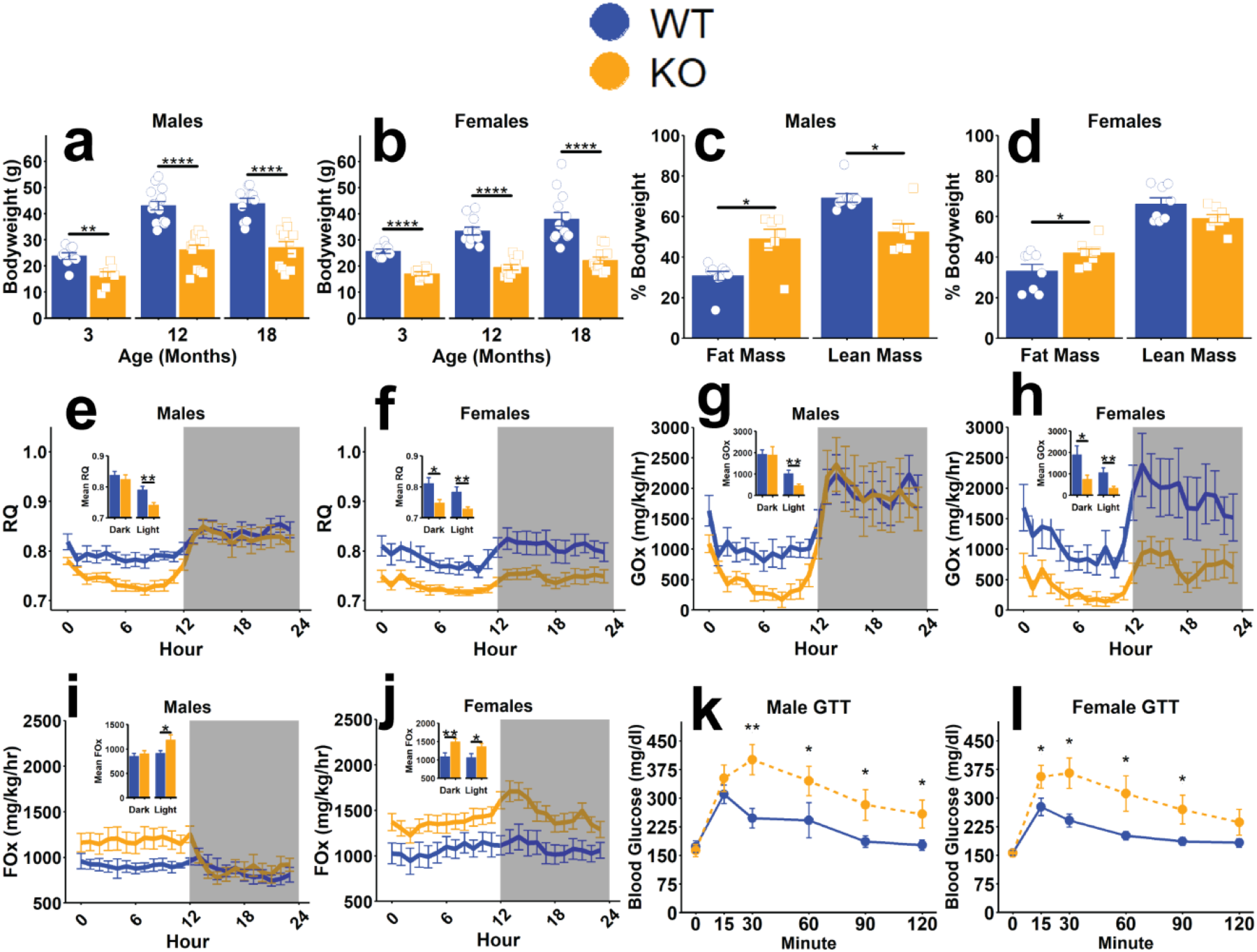
Physiological features of mice with isolated GH deficiency. Bodyweight of male (a) and female (b) mice at the indicated ages. Fat and lean mass as percentages of total bodyweight in male (c) and female (d) mice. 24-hour respiratory quotient (RQ; VCO_2_/VO_2_) measured during indirect calorimetry in male (e) and female (f) mice, with insets representing mean RQ during the dark or light phases as indicated. 24-hour glucose oxidation (GOx) normalized to bodyweight in male (g) and female (h) mice with insets representing mean GOx during dark or light phases as indicated. 24-hour fat oxidation (FOx) normalized to bodyweight in male (i) and female (j) mice with insets representing mean FOx during dark or light phases as indicated. 1g/kg glucose tolerance test in overnight fasted male (k) and female (l) mice. Data presented as mean ± SEM with points representing individual mice. *p<0.05; **p<0.01; ***p<0.001; ****p<0.0001 as determined by two-tailed t-test with the welch correction applied or by Mann-Whitney U-test (c, g, h, k, l). N=7-13 (a-d), 9-15 (e-j), or 9-13 (k, l) per group.

## Discussion

Identification of endocrine signals that control mammalian lifespan is critical for understanding the biology of aging and developing interventions to promote healthy aging. In the present study we demonstrate for the first time, to the best of our knowledge, that targeted disruption of the gene coding for GH extends the lifespan of laboratory mice. In addition to this extended lifespan we also show that these mice are disproportionately fatty despite being lighter than littermate controls, are hypoactive and hypometabolic, rely heavily on fatty acid oxidation to meet their energy demands, and display impairments in glucose metabolism when compared to controls.

It is noteworthy that while the differences in lifespan we observed between KO and WT mice were significant, they are lesser in magnitude than the 40+% extensions reported in other models of somatotrophic disruption (Bartke et al., 2001; Bonkowski et al., 2006; Brown-Borg et al., 1996; Flurkey et al., 2001; List et al., 2011; Sun et al., 2013). While our median KO mouse lifespan of 2.60 years is consistent with the 2.55 (Sun et al., 2013) year and the 2.75 (Sun et al., 2017) year median lifespans observed by us in other models of somatotrophic interruption, it is noteworthy that our median female WT survival of 2.37 years is considerably longer than in these previous reports and may have reduced the magnitude of difference in median survival between KO and WT mice. The 38% difference in median lifespan between our shortest- and longest-lived groups in this study, while certainly an extension, remains below the previously described 40% enhancement. This suggests that while GH deficiency clearly contributes to lifespan extension, an additive effect of additional gene/hormone deficiencies on lifespan may also exist. Previous work suggests that thyroxine treatment, a thyroid hormone secreted in response to thyroid-stimulating hormone, normalizes growth rate and energy metabolism in Ames dwarf mice (Darcy et al., 2016; Panici et al., 2010). While no difference in lifespan was reported following short-term thyroxine treatment (Panici et al., 2010) and observed metabolic effects were transient (Darcy et al., 2016), these reports indicate that the hormone deficiencies apart from GH play relevant roles in producing the dwarf mouse phenotype. Future work examining the contribution of hormones besides GH to the lifespan of mice with an interrupted somatotrophic axis is warranted to understand this powerful mediator of longevity more completely.

A reduced dependance on glucose metabolism to meet organismal energy demands has been consistently reported in various models of somatotrophic interruption(Brooks et al., 2007; Icyuz et al., 2020; Westbrook et al., 2009). The commonality of this observation may indicate that this is an important component in producing the healthy aging physiology of these mice. In the present study we extend these findings to mice with a targeted disruption of GH, suggesting the absence of GH is responsible for these observations. In support of this, GH replacement in dwarf mice normalizes the reduced respiratory quotient observed in these mice (Sun et al., 2017), suggesting increased glucose utilization. As defects in carbohydrate metabolism have been implicated in several age-associated diseases (Brewer et al., 2016), a lower dependency on glucose metabolism may provide resistance to these conditions in mice with interruptions in the somatotrophic axis. While enhanced insulin sensitivity generally accompanies the metabolic changes observed in somatotrophic-interrupted mice (Liu et al., 2004; Wiesenborn et al., 2014; Zhang et al., 2020), here we only detected this in male GH knockout mice. This conflicts with a previous study in this model where insulin sensitivity was increased in both males and females (List et al., 2019). The different genetic backgrounds and ages of mice used between this previous report and our study may contribute to this discrepancy, as mouse genetic background plays a major role in insulin sensitivity (Parks et al., 2015).

In summary, we show for the first time that isolated GH deficiency increases both average and maximal lifespan and that these mice are defined by disproportionate adiposity and impaired glucose metabolism. These data demonstrate that GH itself is a potent regulator of mammalian longevity.

## Methods

### Mice

Mice with a deletion of the GH genomic sequence generated by Regeneron Pharmaceuticals using the VelociGene (Valenzuela et al., 2003) strategy were obtained as live mice from the KOMP Repository (www.KOMP.org) at the University of California, Davis (Stock #: 047836-UCD). These mice have previously been characterized as lacking circulating GH (List et al., 2019) and are referred to as “KO” mice. Mice were housed at up to 7 individuals per cage in a specific pathogen-free facility maintained on a standard 12-h light and 12-h dark cycle at 20–23°C. Original mice were obtained on the C57BL/6N genetic background. C57BL/6N males carrying the mutation were mated with C57BL/6J WT (JAX stock #000664) and with BALB/cByJ WT (JAX stock #001026) females. The offspring of the C57BL/6N x C57BL/6J mating were bred with the offspring of the C57BL/6N x BALB/cByJ mating to produce mice on a mixed C57BL/6N x C57BL/6J x BALB/cByJ genetic background. This was done, as our group previously has (Icyuz et al., 2020; Lasher & Sun, 2023; Nagarajan et al., 2024; Zhang et al., 2020), to increase genetic diversity, increase fecundity, and reduce artifactual findings that arise from utilizing mice on a homogenous genetic background. These mixed-background mice were used for all experiments. All mice had *ad-libitum* access to standard rodent chow (NIH-31 Rat and Mouse diet, 18% protein, 4% fat) and drinking water. All experimental protocols utilizing live animals were approved by the University of Alabama at Birmingham institutional animal care and use committee.

### Metabolic Studies

All mice were 18 months old at the time of metabolic assessment. Body composition was assessed by quantitative magnetic resonance using an EchoMRI body composition analyzer (EchoMRI, Houston, TX). Data collected included body fat mass and lean body mass, which were normalized to total body weight to account for the dramatic differences in body size of our WT and KO mice. Body composition assessment was carried out by the University of Alabama at Birmingham Small Animal Phenotyping Core Facility.

Nutrient utilization and metabolic rate were assessed by indirect calorimetry. Mice were individually housed in comprehensive lab animal monitoring system (CLAMS; Columbus Instruments, Columbus, OH) chambers. This system continuously collects oxygen consumption (VO_2_) and carbon dioxide production (VCO_2_) data every 9 min for each individually housed mouse. This system also monitors mouse activity and food consumption by detecting beam breaks in an infrared laser grid within each chamber and food weight changes in the feeder-balance systems of each chamber. For assessment of mouse activity, only laser beam breaks that were different from the previously broken beam were considered, as this represents movement rather than repetitive stationary activity (i.e. grooming). Mice were individually housed in calorimetry chambers for a period of 48 hours, the first 24 of which were considered acclimation and were not included in data analysis. Respiratory quotient, glucose oxidation, and fat oxidation were calculated as VCO_2_/VO_2_, 4.57(VCO_2_)–3.23(VO_2_), and 1.69(VO_2_)–1.69(VCO_2_), respectively as previously described (Lasher & Sun, 2023; Nagarajan et al., 2024; Simonson & DeFronzo, 1990), and were normalized to body weight. Energy expenditure was calculated as VO_2_(3.815 + 1.232(VCO_2_/VO_2_)) according to the work of Graham Lusk (1928). ANCOVA, with bodyweight as a covariate, was used to compare mean energy expenditure (and VO_2_) between WT and KO groups to properly account for the effect bodyweight has on this metric(Tschöp et al., 2011).

Glucose tolerance tests (GTTs) and insulin tolerance tests (ITTs) were carried out in 16-hour fasted mice (for GTT) or in ambient-fed mice with food removed immediately before testing (for ITT). Blood glucose was taken immediately prior (“minute 0”) and at the indicated time points following an intraperitoneal injection of 1g/kg glucose (for GTT) or 0.7U/kg Humulin-R (Eli-Lilly, Indianapolis, IN, for ITT) prepared in 0.9% saline. A PRESTO handheld glucometer (AgaMatrix, Salem, NH) was used to measure blood glucose.

### Statistical analysis

The Log-rank test and Cox proportional hazard testing were employed to assess overall survival between groups. To compare maximal lifespan, a quantile regression assessment described by Wang and colleagues (Wang et al., 2004) where the proportion of mice alive at the 75^th^ and 90^th^ percentiles of survival were compared. Group means were compared using the unpaired two-tailed t-test with the welch correction applied, the Mann-Whitney U-test (where data were non-parametric), or by factorial repeated measure ANOVA as indicated in the figure legends. To assess energy expenditure and VO_2_, ANCOVA with body weight treated as a covariate was used to control for the effect of size on metabolic rate. For all statistical tests significance was established at *p* < 0.05. Analyses were carried out and figures were generated using the R programming language.

## Acknowledgements

The authors would like to recognize the critical feedback and insightful comments from all members of the Sun Lab that helped conceive this manuscript.

## Funding Information

The authors received research support from the National Institutes of Health AG082327and AG057734 to LYS. The UAB Small Animal Phenotyping Core supported by the NIH Nutrition & Obesity Research Center P30DK056336, Diabetes Research Center P30DK079626 and the UAB Nathan Shock Center P30AG050886A.

## Contributions

ATL took the lead in writing the manuscript. ATL and LYS designed experiments and analyzed data. KL collected data. KL and MF maintained the mouse colony. LYS conceived the study, secured funding, and supervised overall direction. All authors contributed critical feedback that shaped the study and manuscript.

## Conflict of Interest

All contributing authors declare no conflict of interest.

## Data Availability Statement

All data used to generate the statistical analyses and figures in this manuscript are available from the corresponding author upon reasonable request.

## Notes

### Competing Interest Statement

The authors have declared no competing interest.

