## Supplemental Figure 1 for "Isolating the Direct Effects of Growth Hormone on Lifespan and Metabolism"

### Supplemental Figure Legends

**Supplemental figure 1**. Cumulative food intake (a, b) and home cage activity (c, d) recorded during indirect calorimetry in male and female mice as indicated. Oxygen consumption (VO2) in males (e) and ANCOVA analysis for mean VO2 measurements, with bodyweight as a covariate, recorded during indirect calorimetry experiments in males (f). VO2 in females (g) and ANCOVA analysis for mean VO2 measurements, with bodyweight as a covariate, recorded during indirect calorimetry experiments in females (h). Energy expenditure (EE) calculated during indirect calorimetry in males (i) and ANCOVA analysis for mean EE measurements, with bodyweight as a covariate, recorded during indirect calorimetry experiments in males (j). EE calculated during indirect calorimetry in females (k) and ANCOVA analysis for mean EE measurements, with bodyweight as a covariate, recorded during indirect calorimetry experiments in females (l). 0.7 U/kg insulin tolerance test in males (m) and females (n). Data presented as mean ± SEM with points representing individual mice. P-values determined by two-way repeated measure ANOVA (a-d), ANCOVA (f, h, j, l), or Mann-Whitney U-test (m, n). N=9-15 per group.

### Supplemental figure 1

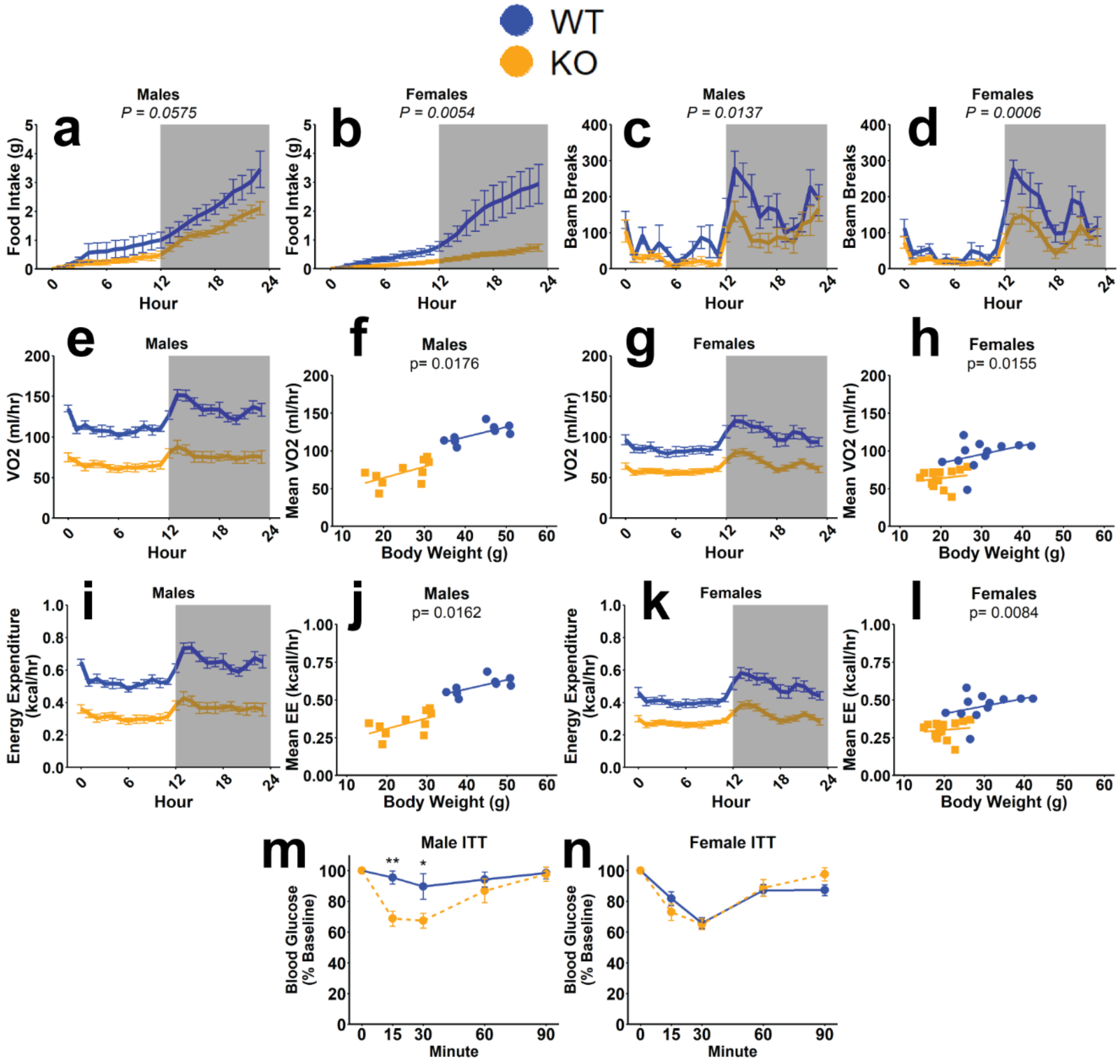
